# SREBP1 activation contributes to fatty acid accumulations in necroptosis

**DOI:** 10.1101/2022.08.22.504843

**Authors:** Daniel Lu, Laura R. Parisi, Omer Gokcumen, G. Ekin Atilla-Gokcumen

**Affiliations:** Department of Chemistry, University at Buffalo, The State University of New York (SUNY), Buffalo, NY 14260, USA; Department of Biological Sciences, University at Buffalo, The State University of New York (SUNY), Buffalo, NY 14260, USA

## Abstract

Necroptosis is a type of programmed cell death. It is characterized by membrane permeabilization and is associated with a strong inflammatory response due to the release of intracellular components due to compromised membrane integrity. We recently showed that the accumulation of very long chain fatty acids (VLCFAs) contributes to membrane permeabilization during necroptosis. However, the mechanisms that result in the accumulation of these cytotoxic lipids remain unknown. Using comparative transcriptomics, we found that sterol regulatory element-binding protein 1 (SREBP1) is activated and that its downstream gene targets result in the accumulation of VLCFAs during necroptosis. We demonstrated that activation of SREBP1 during necroptosis exacerbates cell death. On the contrary, inactivation of SREBP1 reversed the accumulation of VLCFAs, and restored cell death and membrane permeabilization during necroptosis. Collectively, our results highlight a role for SREBP1 in regulating lipid changes during necroptosis and suggest SREBP1 as a potential target for therapeutics to reduce membrane permeabilization during necroptosis.

## INTRODUCTION

Lipids have a plethora of cellular functions. They provide structural integrity for biological membranes^1^ and actively participate in cellular processes, including cell death and survival.^2^ At the intersection of their structural and signaling roles, lipids participate in membrane-related transformations that occur during different cellular processes.^3-5^ For instance, cells can modulate lipid production to provide key metabolites needed for sustained proliferation rates^6,7^ and suppress the biosynthesis of cell survival-related lipids.^8^ Despite these key roles in survival, dysregulated accumulation of lipids can trigger lipotoxicity which contributes to many cellular fates.^9, 10^ In many pathologies, the increased lipid uptake or imbalance of cellular lipolytic and lipogenic activities lead to the accumulation of lipids that cause organelle and membrane damage, perturb fatty acid breakdown, and eventually lead to cell death.^11, 12^ One of the major mechanisms that regulate *de novo* lipid biosynthesis is the transcriptional activation of sterol regulatory elementary binding proteins (SREBPs). SREBP1 and SREBP2, encoded by two different genes, regulate the production of fatty acids and sterols, respectively.^13^

Necroptosis, also known as programmed necrosis, is a form of non-apoptotic cell death.^14, 15^ It is associated with many diseases, including ischemia/reperfusion injuries^16^, myocardial infarction17, and cancer.^18^ Importantly, necroptotic cell death contributes to the inflammatory responses associated with these diseases due to the permeabilization of the plasma membrane upon activation of the death pathway.^19^ The permeabilization of the plasma membrane, in particular, results in the release of cellular content, including cytokines, to the extracellular matrix, which induces an inflammatory response at the organismal level.^20, 21^ Blocking necroptosis has provided encouraging results in reducing inflammation in psoriasis22 and delaying the loss of motor function in a model of Huntington’s disease.^23^ In cancer therapeutics, necroptosis is often viewed as a double edged sword. While functioning as a fail-safe cell death mechanism in apoptosis-resistant cancer cell lines has been shown as a promising strategy, the increase of downstream inflammation and other immune response during necroptosis plays a paradoxical role in tumor progression.^24^ Upon membrane permeabilization, intracellular components including pro-inflammatory cytokines, chemokines, and other growth factors are released into the microenvironment, promoting tumorigenesis.^25^ It has been reported necroptosis promotes the production of pro-inflammatory cytokines^26^ and cancer metastasis in the tumor microenvironment.^27^ Targeting necroptosis, specifically reducing the release of pro-inflammatory stimulants via membrane permeabilization has become a potential therapeutic strategy in combating tumor progression. For instance, a study has demonstrated that the suppression of necroptosis via small molecule inhibitor necrostatin-1 successfully reduces tumor growth in the colitis-associated tumorigenesis mouse model.^28^ Another study has shown inhibition of necroptosis either by small molecule inhibitors or deletion of key proteins in necroptosis activation reduced cell-induced endothelial necroptosis and metastasis.^29^As such cancer progression and metastasis may be linked to the downstream release of inflammatory substances as a result of membrane permeabilization during necroptosis, Thus, specific targets that contribute to membrane permeabilization and toxicity during necroptosis are an exciting frontier for new therapeutics.

The molecular interactions that disintegrate the plasma membrane during necroptosis are well-studied. The membrane modeling that eventually results in membrane rupture is dependent on the activity of Receptor Interacting Protein Kinases 1 and 3 (RIPK1/RIPK3).^14, 30^ Upon activation of death pathways, a complex consisting of RIPK1, RIPK3, and Mixed Lineage Kinase Domain-like protein (MLKL) forms in necroptosis.^31^ This complex phosphorylates MLKL and facilitates the oligomerization of MLKL and its subsequent translocation to the plasma membrane, which compromises the plasma membrane integrity.^32^ Translocation of phosphoMLKL (pMLKL) oligomers to the plasma membrane is primarily driven by electrostatic interactions between the oligomer interface and negatively charged phosphatidylinositol phosphate (PIP)-rich membrane domains^33^. In parallel, recent studies from our group have demonstrated that *S*-fatty acylation of pMLKL by very-long-chain-fatty acids (VLCFAs, fatty acids > 20 carbons^34^) help membrane binding of pMLKL and contribute to membrane permeabilization during necroptosis.^35^ Overall, these two chemical interactions, electrostatic and acylation-mediated membrane binding, cause pMLKL to locate in the plasma membrane and result in the consequent pore formation, which is the basis for the inflammatory nature of necroptotic cell death.^32, 33^ Despite such critical involvement of lipids and the role of membranes in necroptosis, regulatory mechanisms that govern lipid changes in necroptosis are unknown.

In our recent studies toward a better understanding of the contribution of lipids to membrane permeabilization during necroptosis, we found that lipids accumulated during this process.^36^ Specifically, we found that VLCFAs showed profound accumulations at the transcriptional level during this process via the activation of fatty acid synthase (*FASN*) and the elongation of very long chain fatty acids proteins 1 and 7 (*ELOVL1* and *ELOVL7*). We further showed that pMLKL and MLKL, key signaling proteins of necroptosis, undergo fatty acylation by VLCFAs, which improve their membrane recruitment and binding and contribute to plasma membrane permeabilization during necroptosis.^35, 37^ In this work, we investigated the mechanisms of lipid production in this process. Using transcriptomics, we found that there was an overall increase in the expression of enzymes that are involved in *de novo* lipogenesis during necroptosis. Analysis of these changes suggested SREBP1 as a potential mediator of VLCFA accumulation. We showed that SREBP1 was activated and that SREBP1 gene targets that are responsible for the accumulation of VLCFA were upregulated during necroptosis. Further activation of SREBP1 exacerbates cell death whereas inactivating SREBP1 reduced cell death and significantly restored membrane permeabilization in this process. Overall, our results provide a better understanding of the mechanism of necroptosis and pave the way to prioritize lipid-related pathways for therapeutic strategies to delay membrane permeabilization in this process.

## RESULTS

### Transcriptomic analysis highlights the upregulation of lipid biosynthesis-related genes

To investigate the pathways that can be responsible for the transcriptional activation of *de novo* VLCFA biosynthesis during necroptosis, we compared the transcriptomes of necroptotic and control HT-29 cells. Briefly, we extracted RNA from necroptotic and control cells (n= 3 for each condition) and analyzed the mRNA content using Illumina sequencing (see Methods section for details). We then identified genes that were differentially expressed during necroptosis (**Table S1**, *p*_*adjusted*_ < 0.05). **Figure 1A** represents 3,174 genes (1631 upregulated and 1543 downregulated in necroptotic conditions) that showed significant differences between control and necroptotic cells. After manual examination of these genes, we found that 113 of these were lipid-related (**Figure 1B, Table S2**). We validated these findings by measuring the changes in gene expression in an independent set of samples using digital droplet PCR (**Figure S1**, see **Table S3** for primer sequences). Among the lipid-related transcripts that were regulated during necroptosis, we noticed several major targets of SREBP1 and 2. These included key enzymes that regulate *de novo* fatty acid (e.g., fatty acid synthase, *FASN*; elongation of very long chain fatty acids protein 1, *ELOVL1*) ^38, 39^ and cholesterol biosynthesis (e.g., 3-hydroxy-3 methylglutaryl-CoA reductase, *HMGCR*; squalene epoxidase, *SQLE*)^38^ (**Figure 1C**). These observations prompted us to investigate a potential role of SREBPs in regulating lipid composition during necroptosis.

**Figure 1.**
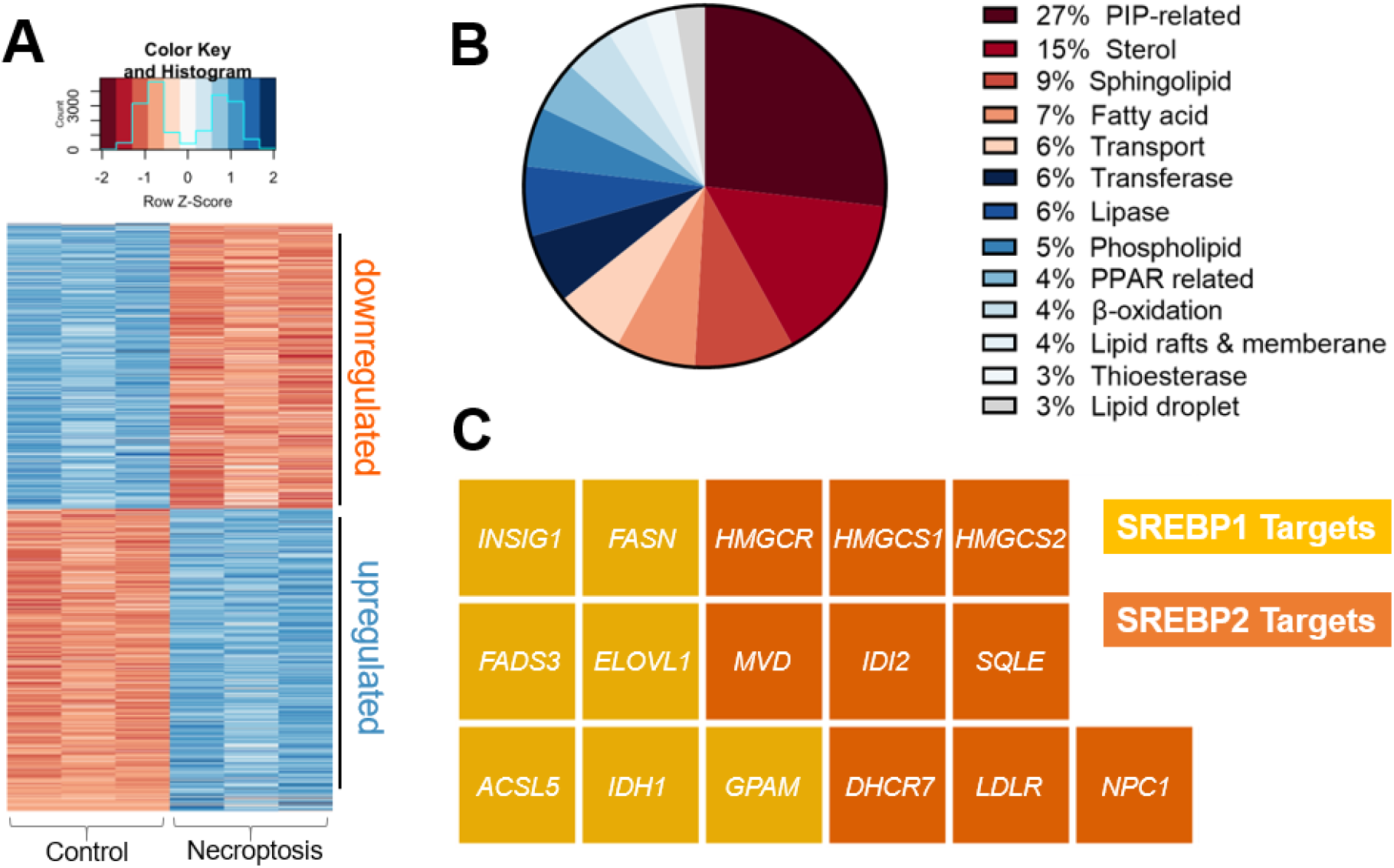
Transcriptomics analysis during necroptosis. (**A**) Heatmap representation of transcripts that are differentially regulated during necroptosis (*p*_adjusted_ < 0.05) (**B**) Among the differentially regulated transcripts, 112 were lipid production, processing, and transport-related. The pie chart shows the distribution of these 112 transcripts based on their function. (**C**) Waffle plot of differentially regulated SREBP targets during necroptosis. Genes of interest were normalized based on the gene expression level in control cells and the fold change was calculated based on the ratio between the average of necroptotic cells with respect to control cells. Genes abbreviation: Insulin induced gene 1, *INSIG1*; fatty acid synthase, *FASN*; fatty acid desaturase 3, FADS3; ELOVL fatty acid elongase 1, *ELOVl1*; acyl-CoA synthetase long-chain family member 5, *ACSL5*; isocitrate dehydrogenase 1, *IDH1*; glycerol-3-phosphate acyltransferase, *GPAM*; isopentenyl-diphosphate delta isomerase 2, *IDI2*; low density lipoprotein receptor, *LDLR*; 3-hydroxy-3-methylglutaryl-CoA synthase 1, *HMGCS1*; Niemann-Pick disease, type C1, *NPC1*; squalene epoxidase, *SQLE*; mevalonate (diphospho) decarboxylase, *MVD*; 7-dehydrocholesterol reductase, *DHCR7*; 3-hydroxy-3-methylglutaryl-CoA synthase 2, *HMGCS2*; 3-hydroxy-3-methylglutaryl-CoA reductase, *HMGCR*.

### SREBP1 is activated during necroptosis

Transcriptional regulation of *de novo* lipid biosynthesis by SREBP1 and 2 are one of the major mechanisms that control lipid production in cells.^38^ SREBP1 primarily activates the transcription of genes that are mainly responsible for the biosynthesis of fatty acids.^40^ SREBP2, on the other hand, activates genes responsible for cholesterol biosynthesis and uptake.^13^ Because we were interested in the accumulation of VLCFAs during necroptosis, we focused on the activation of SREBP1. To investigate the activation of SREBP1 and the effect of potential activation on lipid targets during necroptosis, we first studied its proteolytic maturation using Western blotting. HT-29 cells were pretreated with BV6 and zVAD-fmk for 30 minutes and induced with necroptosis via the addition of TNF-α for 1 h, 3 h, or 5 h. Using western blotting we observed a time-dependent increase in mature SREBP1 levels (up to 3.5 fold, *p* < 0.001) during necroptosis (**Figure 2A-B**). Next, to investigate the effect of SREBP1 maturation on target genes, we used digital droplet PCR and quantified the changes in *FASN* and *ELOVL7* expression, representative SREBP1 targets that control fatty acid synthesis, as necroptosis progressed. Similar to our observation of the time-dependent activation of SREBP1 during necroptosis, the levels of *FASN* and *ELOVL7* increased up to 4-fold during necroptosis (**Figure 2C**). These results show that SREBP1 is activated during necroptosis.

**Figure 2.**
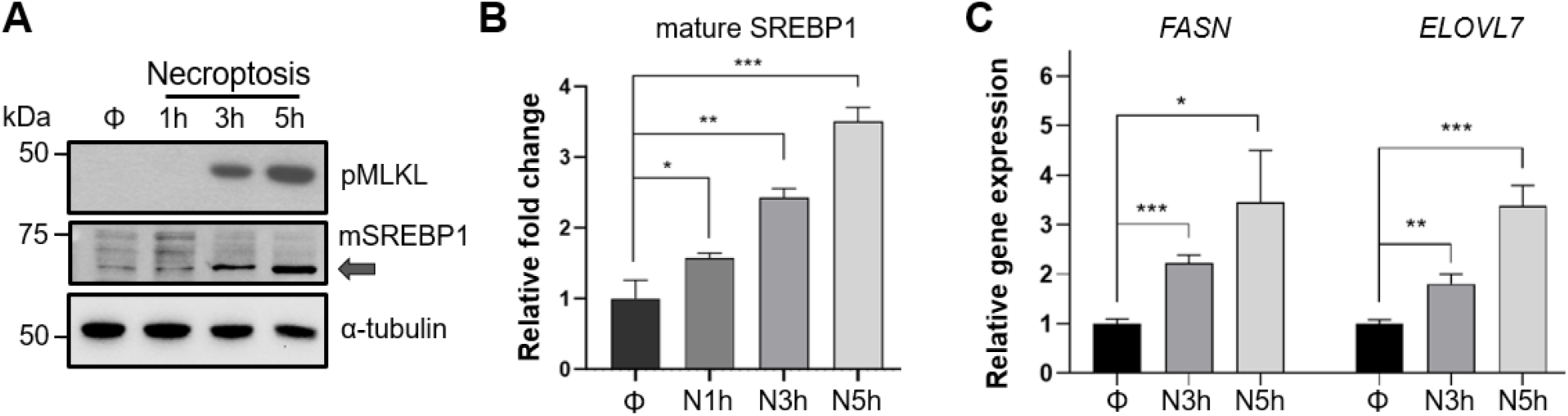
SREPB1 is activated during necroptosis. (**A-B**) Western blot analysis for the time-dependent increase in SREBP1 activation during necroptosis. Cells were induced with necroptosis for 1 h, 3 h, and 5 h. The whole lysate samples were prepared and analyzed. Transcriptional active fragment, mature SREBP1 (mSREBP1) increases time-dependently compared to control during necroptosis. (**A**) A representative Western Blot is shown. (**B**) Quantification of mSREBP1 band intensities shows a significant increase during necroptosis compared to control cells. Φ represents DMSO control, N1h represents 1h necroptosis, N3h represents 3h necroptosis and N5h represents 5h necroptosis. Data represent mean ± 1 SD; n= 3. * represents *p* < 0.05, ** represents *p* < 0.01, *** represents *p* < 0.001. (**C**) SREBP1 target genes showed upregulation during necroptosis. Fold changes in expression of *FASN* and *ELOVL7*, are calculated as the ratio of relative expression of each gene compared with *HPRT1* in necroptotic and control cells. Data represent mean ± 1 SD; n= 3. * represents *p* < 0.05, ** represents *p* < 0.01, *** represents *p* < 0.001.

We note that we also observed similar activation of SREBP2 during necroptosis and increases in expression of target its target genes (**Figures S2A** and **S2B**), suggesting transcriptional activation of cholesterol biosynthetic pathway during necroptosis. To investigate if SREBP2 activation could be involved in lipid regulation during necroptosis, we measured cellular cholesterol levels in control and necroptotic cells using LC-MS as we described earlier.^36^ There was no appreciable change in cholesterol levels during necroptosis (**Figure S2C)**, suggesting that SREBP2 activation does not affect cholesterol pools in necroptosis. Further, we used several small molecule inhibitors of the cholesterol biosynthetic pathway (**Figure S2D**) and investigated the effect of these inhibitors on cell viability during necroptosis. Based on the dose-response curves we generated (data not shown), we chose the highest concentration for each inhibitor that had no significant effect on cell viability on their own and measured cell viability during necroptosis in the presence of these inhibitors. None of the inhibitors we tested had a significant effect on cell death during necroptosis (**Figure S2D**), suggesting that activation of SREBP2 does not affect cell death during necroptosis.

### SREBP1 activation exacerbates necroptosis via the increase in downstream VLCFAs production

To study the functional involvement of SREBP1 activation during necroptosis, we used a pharmacological approach. We used U18666A, an SREBP activator^41^ to investigate whether further activation of SREBP1 via U18666A during necroptosis would lead to increase in VLCFA production and hence increase in toxicity. We pre-treated HT-29 cells with 3 μM U18666A for 24h, induced necroptosis and assessed the levels of SREBP1 activation after U18666A treatment during necroptosis using Western blotting. As we expected, U18666A treatment in necroptosis activated SREBP1 as indicated by the increase in mature SREBP1 levels during necroptosis (**Figure 3A**, left panel is a representative Western blot, right panel is the quantification of mature SREBP1 levels). We then conducted LC-MS targeted analysis to investigate if fatty acids, specifically VLCFAs were also increasing in the presence of U18666A during necroptosis. We extracted lipids from necroptotic and U18666A-treated necroptotic cells and analyzed these extracts as we described earlier.^36^ We observed a ∼3-fold increase in the levels of fatty acids, including VLCFAs that cause membrane permeabilization during necroptosis (**Figure 3B**). The VLCFA accumulation correlates to further activation of SREBP1 observed, potentially exacerbating cell death during necroptosis. To evaluate if the accumulation of VLCFAs could be translated to an increase in necroptosis cell death, we measured the cell viability of cells pretreated with U18666A in necroptosis and indeed we observed a significant decrease in cell viability compared to necroptosis alone (**Figure 3C**). Altogether, these results support that the activation of SREBP1 exacerbates necroptosis.

**Figure 3.**
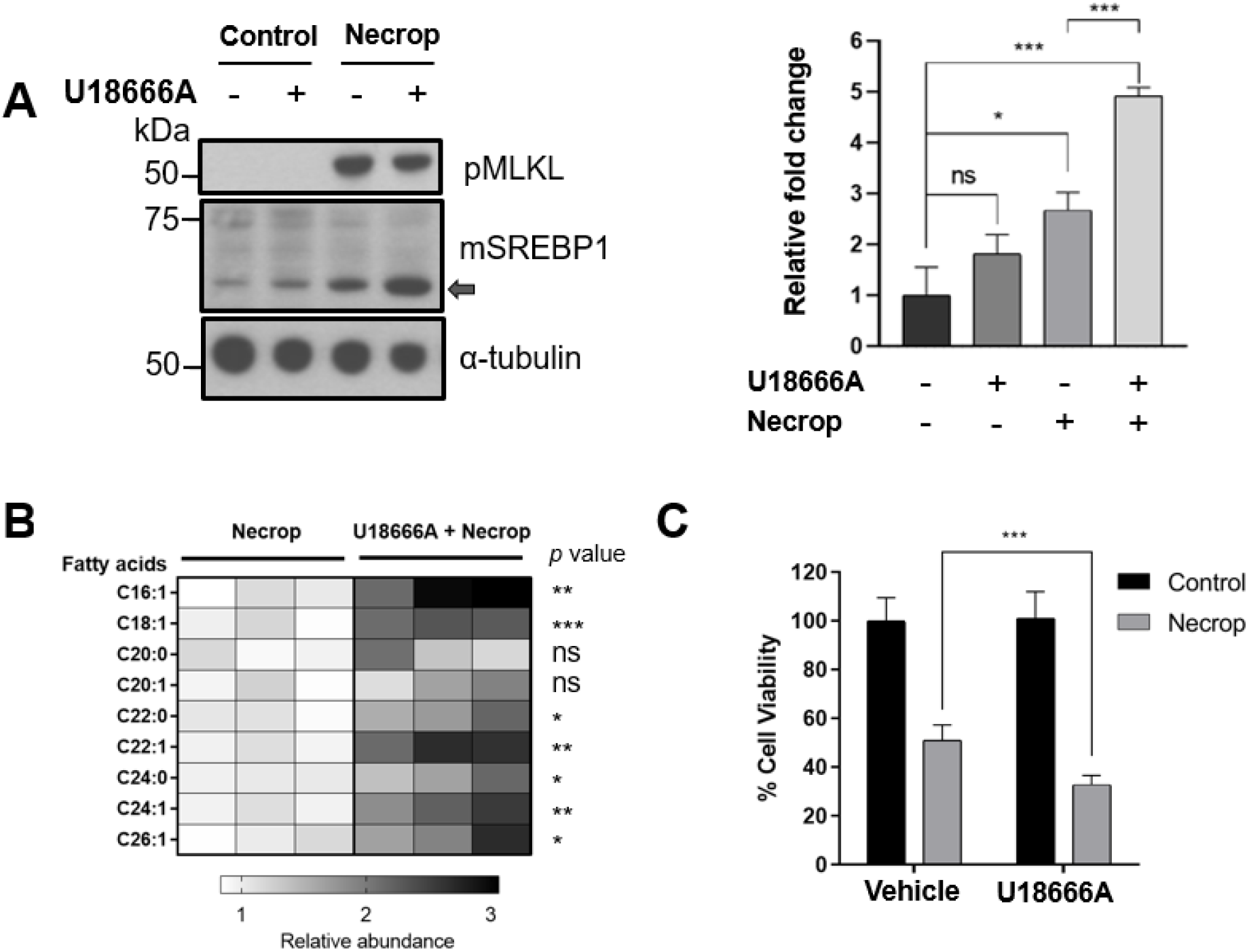
U18666A, an SREBP1 activator contributes to further VLCFA accumulation resulting in an increase in toxicity during necroptosis. (**A**) Western blotting showed SREBPs are further activated after U18666A pretreatment during necroptosis. HT-29 was pretreated with 3 μM of U18666A for 24 h, induced with necroptosis for 3 h. The cells were collected and prepared for western blotting experiment as whole lysate sample. The panel on the right is the quantification of mSREBP1 band intensities which shows a significant increase when pretreated with U18666A during necroptosis compared to control cells and necroptotic cells. Data represent mean ± 1 SD; n= 3. * represents *p* < 0.05, *** represents *p* < 0.001. (**B**) LC-MS showed further VLCFA accumulation in U18666A treated cells compared to necroptosis only cells. HT-29 was pretreated with 3 μM of U18666A for 24 h, induced with necroptosis for 3 h. The cells were collected and were subjected to lipid extraction and LC-MS analysis. n= 3, ns represents *p* > 0.05, * represents *p* < 0.05, ** represents *p* < 0.01, *** represents *p* < 0.001. (**C**) Cell viability during necroptosis decreases when SREBP1 is further activated using U18666A. HT-29 were pretreated with 3 μM of U18666A for 24 h, induced with necroptosis for 3 h, and subjected to MTT cell viability assay. Data represent mean ± 1 SD; n ≤ 5, *** represents *p* < 0.001.

### Preventing SREBP1 activation ameliorates membrane permeabilization and rescues cells from necroptosis

Our results suggest that SREBP1 activation is responsible for the accumulation of VLCFAs that contribute to membrane permeability during necroptosis. We hypothesized that if this is the case, SREBP1 inhibition should then ameliorate cell death in necroptosis. To test this, we first used betulin^42-44^ which results in the degradation of SREBP1, hence reducing overall SREBP1 activity. We pretreated cells with betulin, induced necroptosis, and measured cell viability and plasma membrane permeability during necroptosis. Betulin treatment resulted in a significant rescue from cell death during necroptosis (85% viability in betulin-treated necroptotic cells vs 50% viability in necroptotic cells, **Figure 4A**) and restored plasma membrane permeability, measured by ∼40% decrease in propidium iodide uptake (p < 0.05, **Figure 4B**), supporting our hypothesis that SREBP inhibition reduces cell death during necroptosis.

**Figure 4.**
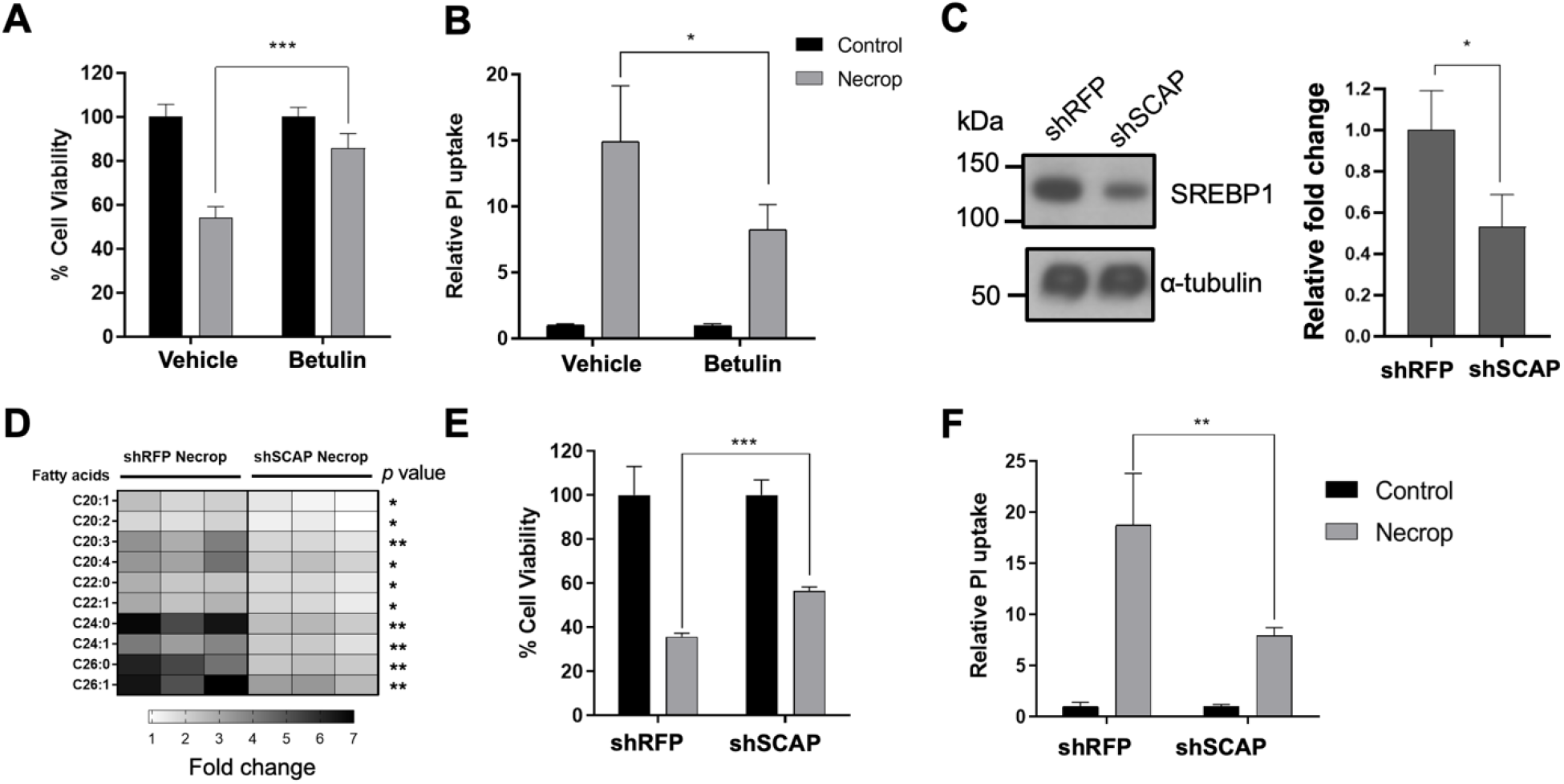
Betulin, a SREBP1 deactivator, and shSCAP knockdown rescue cells from membrane permeabilization. (**A**) Cell viability during necroptosis increases when SREBP1 is deactivated by small molecule inhibitor betulin. HT-29 was pretreated with 2 μM of betulin for 24 h, induced with necroptosis for 3 h, and subjected to MTT cell viability assay. Cell viability was normalized to the corresponding control condition. Data represent mean ± 1 SD; n= 5, *** represents *p* < 0.001. (**B**) Betulin prevents membrane permeabilization in necroptosis as indicated by reduced fluorescence intensity in propidium iodide uptake. HT-29 was pretreated with 2 μM of betulin for 24 h, induced with necroptosis for 3 h, and subjected to propidium iodide uptake experiment. Data represent mean ± 1 SD; n= 5, * represents *p* < 0.05. (**C**) SREBP1 precursor depletion in shSCAP knockdown cells. Red fluorescent protein (RFP) is used as a knockdown control. The panel on the right shows the quantification of precursor SREBP1 depletion compared to shRFP cells. (**D**) Targeted LC-MS analysis showed lowered VLCFAs pool in shSCAP cells during necroptosis when compared to shRFP necroptotic cells. The fold changes were calculated by normalizing to the corresponding control condition. n= 3, * represents *p* < 0.05, ** represents *p* < 0.01 (**E**) Cell viability during necroptosis increases in shSCAP cells. shSCAP cells were induced with necroptosis for 3 h and subjected to MTT cell viability assay. Cell viability was normalized to the corresponding control condition. Data represent mean ± 1 SD; n= 5, *** represents *p* < 0.001. (**F**) shSCAP knockdown prevents membrane permeabilization in necroptosis. PI uptake was normalized to the corresponding control condition. The reduction of propidium iodide uptake in shSCAP cells compared to shRFP cells during necroptosis. Data represent mean ± 1 SD; n= 5, ** represents *p* < 0.01.

To validate observed rescue in cell viability and reduction in membrane permeabilization using the small molecule, we used the lentiviral knockdown approach to deactivate SREBP1, targeting SREBP cleavage activating protein (SCAP). SCAP is a chaperone protein that is needed for translocation of precursor SREBP1 from ER to Golgi where proteolytic cleavage and release of the active transcription factor occurs.^13^ We knocked down SCAP in HT-29 cells (**Figure S3A**, shSCAP) and confirmed that precursor SREBP1 levels are depleted in the knockdown model (**Figure 4C**) as well as the target genes of SREBP1 (**Figure S3B**). We then investigated the accumulation of fatty acids and membrane permeabilization in shSCAP cells during necroptosis. shSCAP cells showed a less profound accumulation of fatty acids in necroptosis when compared to control necroptotic cells (shRFP, **Figure 4D**) and a similar rescue in cell viability in necroptosis (**Figure 4E**). Importantly, shSCAP cells exhibited a reduction in propidium iodide (PI) uptake during necroptosis, suggesting a decrease in membrane permeabilization as compared to control necroptotoc cells (**Figure 4F**). Overall, these results strongly support that that inactivation of SREBP1 increases cell viability and restores membrane permeability during necroptosis.

### Potential mechanism of SREBP1 activation during necroptosis

One of the major mechanisms that control the SREBP1 activation is the cholesterol levels at the endoplasmic reticulum (ER).^45^ When cholesterol level at the ER is low, SREBP1 is proteolytically processed to its transcriptionally active form, mSREBP1. However, studies have suggested other mechanisms for the activation of SREBP1. One study showed that SREBP1 can be proteolytically activated by low phosphatidylcholine levels and identified phosphatidate phosphatase-1 as an activator of SREBP1.^46, 47^ We analyzed phosphatidylcholine levels and found that they do not exhibit significant changes during necroptosis (**Figure S4A**). In parallel, our previous results showed no significant changes in phosphatidic acids and diacylglycerol levels during necroptosis^48, 49^; hence phospholipid-dependent activation of SREBP1 is not likely in necroptosis. Another study has shown peroxisome proliferator-activated receptors–α (PPARα) is involved in SREBP1 activation.^50^ Our transcriptomics results show that PPARα was downregulated in necroptosis (**Table S1**, *p*_*adjusted*_ = 0.0005), which does not support PPARα-mediated SREBP1 activation in necroptosis.

Therefore, it is possible that SREBP1 activation in necroptosis can be due to low cholesterol levels at the ER, either due to impaired biosynthesis or trafficking. Since we have shown that cellular cholesterol level does not change and that inhibition of cholesterol biosynthesis does not affect cell viability during necroptosis, we investigated cholesterol localization. We used fluorescence microscopy to visualize intracellular cholesterol localization during necroptosis. We induced necroptosis, stained cells with filipin, a fluorescent polyene molecule that binds to free cholesterols^51^, and wheat germ agglutinin (WGA) as a membrane marker^4, 52^. We then quantified the fluorescence signal at the plasma membrane, based on WGA signal. We observed that in necroptosis, cholesterol showed an increased plasma membrane localization (**Figure S4B-C**). Specifically, in necroptotic cells, filipin intensity was ∼2.5-fold higher at the plasma membrane as compared to control cells (**Figure S4C**, n = 20, p < 0.001). These results suggest that cholesterol localization might be dysregulated during necroptosis and responsible for the activation of SREBP1 during necroptosis. Further investigations are warranted to fully establish the involvement of cholesterol in SREBP1 activation during necroptosis.

## Summary and conclusions

We have previously shown that fatty acid biosynthesis is activated at the transcriptional level during necroptosis and causes the accumulation of VLCFAs during this process. We also showed that VLCFA accumulation contributes to membrane damage-induced cell death in necroptosis. Here, we used transcriptomics, targeted gene expression analysis, and pharmacological perturbations, and found that sterol regulatory element-binding protein 1 (SREBP1) is activated during necroptosis. We showed that using the SREBP1 activator triggers downstream activation of lipid production and increases toxicity during necroptosis. On the contrary, inactivating SREBP1 rescues necroptotic cell death and reduces membrane permeabilization. These results strongly support that SREBP1 is responsible for the accumulation of toxic VLCFAs during necroptosis.

Activation of *de novo* lipogenesis due to SREBP activation is related to several pathologies, including the progression of different forms of cancers.^53, 54^ In these systems, increased *de novo* lipid biosynthesis provides the necessary building blocks for uncontrolled cell division.^53^ Other than potential implications in cancer therapeutics, blocking SREBP activation and decreasing lipid content *in vivo* has shown promising potential to combat metabolic diseases such as type II diabetes and atherosclerosis.^55^ Our results reveal a new role of SREBP1 activation, shed light on important mechanisms of lipid regulations during necroptosis, and provide potential targets for diseases that involve this process.

## Experimental Section

### Materials

HT-29 (human colorectal adenocarcinoma epithelial) cells were obtained from the American Type Culture Collection. Dulbecco’s Modification of Eagle’s Medium (DMEM), penicillin-streptomycin and trypsin were acquired from Corning. Fetal Bovine Serum (FBS) and dimethyl sulfoxide (DMSO) (Cat #42780) were obtained from Sigma-Aldrich and Alfa Aesar, respectively. Antibodies were obtained from Abcam (rabbit monoclonal anti-phospho S358-MLKL, Cat. #ab187091; rabbit polyclonal anti-SREBP1, Cat # ab28481), Cayman (rabbit polyclonal anti-SREBP2, Cat. # 10007663), Millipore Sigma (mouse monoclonal anti-α-tubulin, Cat# T9026) Promega (goat anti-rabbit HRP conjugate, Cat. # W4011), and Jackson Immunoresearch Lab (goat anti-mouse HRP conjugate, Cat. #115-035-174). BV6 (Cat #S7597) and simvastatin (Cat. # S1792) was obtained from Selleck Chemicals, zVAD-FMK was obtained from Enzo (Cat # ALX-260-020), TNF-α was obtained from R&D Systems (Cat #210-TA/CF). Atorvastatin (Cat. # 10493). BIBB 515 (Cat # 10010517), Ro48-8071 (Cat # 10006415), U18666A (Cat. # 10009085), Betulin (Cat#11041), and filipin III (Cat# 70440) were obtained from Cayman Chemical. TAK-475 (Cat. # SML2168) and terbinafine (Cat. # 78628-80-5) were obtained from Sigma. Wheat Germ Agglutinin Alexa Fluor™ 594 Conjugate (Cat# LSW1162) was obtained from Invitrogen. Protease inhibitor (Cat #A32955) were obtained from Thermo Fisher Scientific. MTT reagent (Cat # L11939) was obtained from Alfa Aesar. ProLong™ Gold Antifade Mountant (Cat# P36930) and Propidium iodide (Cat# P1304MP) were obtained from ThermoFisher. E.Z.N.A. HP total RNA kit (Cat# R6812-02) was obtained from OMEGA bio-tek. iScript Advanced cDNA Synthesis Kit (Cat# 172-5037) and ddPCR Supermix for Probes (186-3026) were obtained from Bio-Rad. PCR primers listed in table S3 were obtained from Integrated DNA Technologies.

## Methods

### Cell culture and culture conditions

Human colorectal adenocarcinoma HT-29 cell (ATCC® HTB-38™, adult Caucasian female origin) were cultured at 37°C in 5% CO_2_ atmosphere in DMEM supplemented with 10% (v/v) fetal bovine serum and 1% (v/v) penicillin/streptomycin solution. Cells were cultured for approximately 2 months and were routinely checked for mycoplasma infection.

### Inducing Necroptosis

To induce necroptosis, HT-29 cells were initially sensitized to TNF-dependent cell death pathway by SMAC mimetic BV6 (1 μM), and were co-treated with pan-caspase inhibitor zVAD-FMK (25 μM) and incubated for 30 min at 37 °C. Cells were then treated with TNF-α (10ng/mL) and were incubated for 3 h.

### Cell Viability Assays

HT-29 cells were seeded in 96-well plates with 30,000 cells per well (for 2h pretreatment), 15,000 cells per well (for 24h pretreatment) or 10,000 cells per well (for 48h pretreatment. For shSCAP knockdown viability assay, 30,000 cells per well were seeded. The cells were left to attach for 20 h before pretreatment with small molecule inhibitors or induction of necroptosis. After the designated pretreatment time and/or necroptotic treatment, the 96-well plate was centrifuged for 2 min at 200 rcf at room temperature. The media was removed from each well and replaced with 200 μL of fresh media that contains 5 mg/mL of MTT reagent. The plate was incubated at 37 °C for 2.5 h and then centrifuged for 2 min at room temperature. 155 μL of media was removed from each well and 90 μL of DMSO was added back. The plate was then incubated at 37 °C for 10 min to solubilize the formazan crystal and was centrifuged at 1000 rcf for 2 min at room temperature. Absorbance was measured at 550 nm using Biotek Synergy H1 plate reader. To calculate the percent viability of treated cells compared to control cells, the raw absorbance values of cells were subtracted from the average absorbance values of blanks. For the effect of U18666A on cell viability during necroptosis, corrected absorbance values were normalized to the average absorbance values of vehicle-treated control cells and were expressed as percentage cell viability. For the effect of betulin on cell viability during necroptosis, corrected absorbance values were normalized to the average absorbance values of each condition’s treated control cells and were expressed as percentage cell viability Results are representative of at least two independent experiments, with n≥ 3 each.

### Western blotting

HT-29 cells (1×10^7^) were plated in 10 cm dishes for 20 h attachment. Necroptosis was induced in the cells using the method described above. For experiments with U18666A, 4 × 10^6^ cells were plated in 10 cm dishes and left to attach for 20 h. Cells were pretreated with U18666A (3 μM final concentration) or DMSO for 24 h. For the shSCAP knockdown cells, shRFP and shSCAP (approximately 3×10^6^ cells) were plated and left to attach for 20h. After the cells received the pretreatment and / or induction of necroptosis, cells were scraped from the plates on ice and transferred to 15 mL centrifuge tubes. Plates were rinsed with cold 1x PBS after scraping, which was also transferred to the centrifuge tube. Cells were centrifuged for 5 min at 500 rcf at 4°C. The supernatant was decanted and the cell pellets were washed two more times with cold PBS. The supernatant was decanted and the cell pellet was stored at -80°C. The frozen cell pellets were thawed on ice and were lysed by re-suspending in lysis buffer (1 pellet of protease inhibitor dissolved in 10 mL of Mammalian Protein Extraction Reagent) for 45 mins on ice. The cell lysates were then centrifuged (16,000 rcf, 15 min, 4 °C) and the amount of protein was measured using Bradford Protein Assay according to the manufacturer’s instructions. The samples were normalized based on the lowest protein amount and were diluted with 1:1 with a solution of 95% 5x loading and 5% 2-Mercaptoethanol. The samples were boiled for 10 min and stored in -20 °C

Samples were loaded and separated with sodium dodecyl sulfate polyacrylamide gel (10%) electrophoresis at 150 V. Polyvinyl difluoride (PVDF) membranes were then activated in methanol for five minutes. Once the membrane was activated, the separated proteins were transferred onto the PVDF membranes at 50 V for 2 h. After transfer, the membranes were blocked in 10% non-fat dry milk in tris-buffered saline (TBS)-Tween [10 mM Tris-base, 100 mM NaCl, 0.1% Tween 20 (pH 7.5)] at room temperature for one hour. Membranes were washed three times at 10 minute intervals in TBS-Tween. The corresponding membranes were incubated 1 h at room temperature or overnight at 4°C with primary antibodies (1:500 dilutions for SREBP1, 1:1000 dilution for SREBP2 1:1000 for pMLKL, 1:10000, for α-tubulin). After the designated incubation time, the membranes were washed four times with TBS-Tween for 10 minute each time. The secondary antibodies used were 1:2000 anti-rabbit HRP conjugate and 1:1000 anti-mouse HRP conjugate. Secondary antibodies were diluted with 5% non-fat dry milk in TBS-Tween and incubated for one hour at room temperature. The membranes were then washed again with TBS-Tween three times, 10 min each time, prior to developing with Super Signal West Pico kit (Thermo Scientific).

### Quantification of mSREBP1 bands

Quantitative analysis of Western blot images was performed by using Fiji-ImageJ software. A frame for measurement was developed by using the rectangle tool of Fiji-ImageJ to cover the largest band of the protein of interest. The same frame size was then applied for measuring the intensity of the other protein bands. The background was also measured with the same frame to obtain background intensity measurements. For the background measurement, a region near the protein of interest was used. The measured intensities from the protein of interest and background were then inverted by deducting measurements from 255. The inverted intensities of the protein bands of interest were corrected with the inverted intensities of the background. The relative intensities were obtained by dividing the corrected intensity of mSREBP1 by the corrected intensity of the loading control (α-tubulin, n= 3).

### Lentiviral knockdown of SCAP in HT-29

Knockdown of SCAP in HT-29 cells was performed similarly as described previously 36. Briefly, shRNA (in the pLKO.1-Puro lentiviral vector) targeting SREBP1 and SCAP plasmid DNA was packaged into lentivirus particles in HEK293T cells via co-transfection with psPax2 and pCMV-VSV-G using X-tremeGENE 9 transfection reagent for 48 h. The viral particles were collected and filtered with a 0.45 µM filter. For lentiviral transfection in HT-29 cells, 5 × 105 cells were plated and left to attach overnight. The following day, the media was replaced with 8 µg/mL polybrene media and incubated for 5 min. after 5 min, 50 µL of viral suspension was added to each designated well and incubated for 48 h. The virus-containing media was removed and transfected cells were transferred to a new flask in 2 µg/mL puromycin-containing media. The cells were selected for 2 days and the media was changed to 1 µg/mL puromycin and were then cultured continuously in the 1µg/mL puromycin-containing media.

### Propidium iodide uptake

For betulin pretreatment, HT-29 cells (15,000 cells per well) were seeded in 96-well plates and were left to attach for 20 h. Betulin (final concentration 2 µM) was added for 24h pretreatment, followed by induction of necroptosis for 3h. For shSCAP knockdown cells, shSCAP and shRFP cells (30,000 cells per well) were seeded in 96-well plates and were left to attach for 20 h. Knockdown cells were induced with necroptosis for 3 h. After 3 h necroptosis, the plate was centrifuged for 2 min at 200 rcf at room temperature. Media in the plate was removed and added back with 200 µL propidium iodide (5 µg/mL in PBS). Cells were incubated with propidium iodide for 35 min at 37 °C. After incubation, the plate was centrifuged for 2 min at 300 rcf. The fluorescence intensity was measured using Biotek Synergy H1 microplate reader (excitation wavelength of 535 nm and emission wavelength of 625 nm). Relative fluorescence units were reported as calculated after performing blank subtraction and normalized to vehicle control or corresponding control.

### Gene expression measurements

HT-29 cells (approximately 1.5×10^6^ per well) were seeded onto a 6-well plate and were left to attach for 20h. Cells were induced with necroptosis for 3 h and 5 h or DMSO as control (n = 3). For the shSCAP knockdown cells, shRFP and shSCAP (approximately 1.5×10^6^ cells per well) were plated in 6-well plates (n=3) and left to attach for 20h. After induction of necroptosis or attachment, cells were collected and the total RNA was extracted according to E.Z.N.A HP Total RNA kit manufacturer’s protocol. BioRad’s Droplet Digital PCR: QX200 System was used to convert to cDNA with iSCript cDNA synthesis kit, and to amplify and quantify cDNA. Reaction mixtures including primers, probes, and cDNA were prepared with ddPCR Supermix for Probes (no dUTP) according to the manufacturer’s instructions. An automated droplet generator was used to create water-oil emulsion droplets from the reaction mixtures. A QX200 droplet reader was used to quantify HEX and FAM fluorescence in 20,000 droplets per sample. BioRad Quantasoft software was used for data analysis. The relative expression was calculated based on the ratio between the gene of interest and the housekeeping gene (*HPRT1 or GAPDH*). The foldchange was calculated by normalizing the relative expression in necroptosis to the DMSO control. The primer sequences are given in Table S3.

### Lipid extraction and LC-MS data acquisition for fatty acids and PC analysis

Procedures for lipid extraction and LC-MS methods were adapted from a previous study (Parisi et al., 2017). Cell pellets were thawed on ice and resuspended in 1 mL cold PBS. A 30 μL aliquot of this cell suspension was taken and added to an equal volume of lysis buffer on ice for 1h. The protein concentration of the sample was then determined using Bradford Protein Assay. The remaining 970 µL cell suspension was transferred to a Dounce tissue homogenizer to which 1 mL of cold methanol and 2 mL cold chloroform were added. The mixture was homogenized with 30 strokes and then transferred to a 2-dram glass vial. The vials were centrifuged (500 rcf, 10 min, 4 °C) and the chloroform layer was transferred to a 1-dram glass vial. From the chloroform layer, 1.2 mL was further transferred into a new 1-dram to ensure an equal volume was taken from each sample. The chloroform was removed by rotary evaporation and the dried lipids were stored at -80 °C until ready to be analyzed. Lipids were resuspended in chloroform with volume normalized to protein concentration.

LC-MS analyses were performed using an Agilent Infinity 1260 HPLC/ Agilent 6530 Jet Stream ESI-QToF-MS system. Reverse phase gradient elution chromatography was used for separations. Mobile phase A was composed of 95% water and 5% methanol. Mobile phase B was composed of 60% isopropanol, 35% methanol, and 5% water. For positive ionization mode analyses, 0.1% (v/v) formic acid and 5 mM ammonium formate were added to both mobile phases to improve ionization. For negative ionization mode analyses, 0.1% (w/v) ammonium hydroxide was added instead. LC gradient started with 5 min of 0% B at 0.1 mL/min, then the flow rate was increased to 0.5 mL/min. The mobile phase gradient was ramped from 0% B to 100% B over 60 min, maintained at 100% B for 7 min, then switched to 0% B for 8 min. For positive ionization mode analyses, a Luna C5 column was used as the stationary phase along with a C5 guard cartridge. For negative ionization mode analyses, a Gemini C18 column was used along with a C18 guard cartridge. A DualJSI fitted electrospray ionization (ESI) source was used for MS analysis with a capillary voltage of 3500 V and fragmentor voltage of 175 V. Drying gas temperature was 350 °C with a flow rate of 12 L/min. Data were collected using an m/z range of 50-1700 in extended dynamic mode. Tandem mass spectrometry data were collected using the following collision energies: 15, 35, 55, and 75 eV for each m/z. Abundances were obtained from peak integration of the extracted ion chromatograms. Fatty acids were analyzed in negative mode as[M-H]-adducts, and PC were analyzed in positive mode as [M+H]+ adducts. For fatty acids or phosphatidylcholine that were detected in blank injections, blank subtractions were carried out.

### Analysis of fatty acids in shSCAP during necrotposis

shRFP and shSCAP (8 × 10^6^ cells) were plated in 10 cm dishes for ∼20 h attachment. The cells were added BV6 (1 μM), and zVAD-FMK (25 μM) and incubated for 30 min at 37 °C. Necroptosis was induced using TNF-α (10 ng/ mL final) for 3 h. The cells were then collected on ice, centrifuged, washed with PBS, and then stored at -80 °C until lipid extraction. Lipid extraction and fatty acid analysis in LC-MS were performed as described above. Abundances of different fatty acids were obtained from peak integration and blank subtraction was carried out. Fold changes of fatty acids in shRFP necroptosis were calculated by dividing the abundance of the fatty acid in shRFP necroptosis by the average abundance of shRFP control. Fold changes of fatty acids in shSCAP necroptosis conditions were calculated by dividing the abundance of the fatty acid in shSCAP necroptosis by the average abundance of shSCAP necroptosis.

### Analysis of fatty acids and phosphatidylcholines (PCs) in U18666A treated cells during necroptosis

HT-29 cells (4 × 10^6^ cells) were plated on 10 cm dishes for ∼20 h attachment. Cells were pretreated with U18666A (3 μM final concentration) or DMSO for 24 h. Necroptosis was induced using TNF-α (2 ng/ mL final, with BSA as carrier protein) for 3h. After 3 h, the cells were collected on ice, centrifuged, washed with PBS, and then stored at -80 °C until lipid extraction. Lipid extraction and lipid analysis in LC-MS was performed as described above. Abundances of different fatty acids and PC were obtained from peak integration and blank subtraction was carried out. Relative abundance of the fatty acids in necroptotic cells or U18666A treated necroptotic cells were calculated by dividing the abundances of the lipids by the average abundance of necroptotic cells. Relative abundance of PCs were calculated by dividing the abundances of the lipids by the average abundance of control cells.

### Analysis of cholesterol during necrotposis

HT-29 cells (1 ×10^7^) were plated in 10 cm dishes for ∼20h attachment. The cells were added BV6 (1 μM), and zVAD-FMK (25 μM) and incubated for 30 min at 37 °C. Necroptosis was induced using TNF-α (10 ng/ mL final) for 3 h. The cells were collected and lipids were extracted as described above. Briefly, cell pellets were thawed on ice and resuspended in 1 mL cold PBS. A 30 μL aliquot of this cell suspension was taken for protein measurement using a Bradford Protein Assay. The remaining suspension was transferred to a Dounce tissue homogenizer to which 1 mL of cold methanol and 2 mL cold chloroform were added. The mixture was homogenized with 30 strokes and then transferred to a 2-dram glass vial. The vials were centrifuged (500 rcf, 10 min, 4 °C) and the chloroform layer was transferred to a 1-dram glass vial. The methanol: PBS layer was extracted again by adding another 2 mL of chloroform. The mixture was vortexed for 5s, 3 times, and centrifuged (500 rcf, 10 min, 4 °C). The chloroform layer is transferred into the 1-dram vial, from which 3 mL of chloroform layer was further transferred into a new 1-dram vial. The chloroform layer was dried using Reacti-Vap™ Evaporators. The samples were reconstituted in chloroform with volumes normalized according to protein concentration.

LC-MS analyses were performed similarly as described above. Briefly, the analysis of cholesterol was carried out using an Agilent Infinity 1260 HPLC/ Agilent 6530 Jet Stream ESI-QToF-MS system. 5 μL of the sample was injected for each analysis. Reverse phase gradient elution chromatography was used for separations. Cholesterol detection was carried out in positive ionization mode. Mobile phase A was composed of 95% water and 5% methanol. Mobile phase B was composed of 60% isopropanol, 35% methanol, and 5% water. For positive ionization mode analyses, 0.1% (v/v) formic acid and 5 mM ammonium formate were added to both mobile phases to improve ionization. LC gradient started with 5 min of 0% B at 0.1 mL/min. The gradient was ramped to 100% B over 60 min, and maintained at 100% B for 7 min. For positive ionization mode analyses, a Luna C5 column was used as the stationary phase along with a C5 guard cartridge. A DualJSI fitted electrospray ionization (ESI) source was used for MS analysis with a capillary voltage of 3500 V and fragmentor voltage of 175 V. Drying gas temperature was 350 °C with a flow rate of 12 L/min. The abundance of cholesterol was obtained from peak integration and blank subtraction was carried out. Fold changes were calculated by dividing the abundance of cholesterol in necroptotic cells by the average abundance of control cells.

### Filipin staining for cholesterol localization during necroptosis

HT-29 cells (15 × 10^4^) were seeded onto glass coverslips in 24-well plates and were left to attach for 20 h. Necroptosis was induced using 2 ng/mL TNF-α for 3 h. After induction of necroptosis, the coverslips were washed three times with PBS and the cells were fixed using 3.7% paraformaldehyde for 30 min at room temperature. Coverslips were washed with PBS and the cells were stained with Filipin (1:50 dilution in PBS) overnight at 4 °C. The following day the coverslips were washed three times with PBS, and were stained with WGA-594 (1:2000 dilution in 2% BSA-TBS) for 1 h at room temperature. The coverslips were subjected to three PBS washes and were mounted on glass slide using Prolong Gold Antifade Reagent. Images were acquired on a LSM710 confocal microscope (Carl Zeiss, Oberkochen, Germany). For quantification of filipin intensity localization in control and necroptotic cells, Fiji-imageJ software was used. Quantification of filipin intensity for control and necroptotic cells is done in the following steps: (1)The sum of projection was obtained. A straight line (line width/thickness = 1, line size is 32.38 µm) across the cell was drawn using the line scan feature in Fiji (a total of 10 cells were chosen for quantification) (2) Filipin and WGA-594 intensity profiles were obtained according to the line drawn across the cell. The filipin signal at the plasma membrane was obtained based on the intensity of the plasma membrane marker, Alexa 564 WGA. Cytoplasmic filipin signal was obtained the same way. (3) The ratio of the filipin signal at the plasma membrane and cytoplasm was reported was used to assess cholesterol localization.

### Transcriptomics

Samples were processed by TrueSeq using standard library preparation workflow and single-read sequenced via Illumina HiSeq 2500. We conducted FASQC and MultiQC to test the quality of reads Quality control of paired reads was performed using FastQC and MultiQC.^56^ We used Kallisto to align the reads to Hg19 and quantify expression of each gene (ref).^57^ We then used DESeq2 to compare the transcript abundances for each gene between control and target samples and identify differentially expressed genes using the Benjamini and Hochberg false discovery rate (FDR) corrected adjusted *P*-value, *P* < 0.05.^58^ We provide the gene expression levels and the results of comparative analysis in **Table S1**. We are in the process of uploading the raw RNAseq dataset to Short Read Archive (SRA, https://www.ncbi.nlm.nih.gov/sra).

### Statistics

All statistical analysis for the western blot quantifications, cell viability experiments, LC-MS lipid abundance, and ddPCR were performed using unpaired Student’s *t*-test. *p* values and numbers of replicates in all figures are indicated in figure legends, where *** *p* < 0. 001, ** *p* < 0.01, * *p* < 0.05, and ns is *p* > 0.05. For transcriptomic analysis, the Log_2_ fold changes, *p* and *p*_adjusted_ values (calculated by Wald test for multiple hypotheses) were calculated using DESeq2.

## Supplemental Information

Supplemental Information includes 4 figures, 3 tables.

## Author Contributions

Experiments were designed by D.L., L.R.P., O.G., and G.E.A-G. The experiments were conducted by D.L. and L.R.P. Transcriptomics analysis was done by O.G. The manuscript was written by D.L., L.R.P., O.G., and G.E.A-G. The study was directed by G.E.A-G.

## Acknowledgments

We acknowledge the support from the National Science Foundation grant (MCB1817468 to G.E.A.G.). We thank UB North Campus Imaging Facility for confocal microscopy. We also thank Marie Saitou for her assistance with transcriptomics.

