## Supplementary information for "SREBP1 activation contributes to fatty acid accumulations in necroptosis"

**Table S1 (Excel file). Results of transcriptomic data. Transcripts detected in samples from control and necroptotic cells are reported.**
**This table is provided as a separate spread sheet. The spreadsheet contains two tabs:**
**The first tab ("raw read depths") contains all genes detected in control and necroptotic cells:**
Column A shows the ensemble gene ID for detected transcripts.
Columns B-D show the values of read depth of transcripts in control cells (n= 3)
Columns E-G show the values of read depth of transcripts in necroptotic cells (n= 3)
Column H shows the HUGO gene nomenclature (hgnc_symbol) for identified transcript
Column I shows the description of hgnc_symbol
Column J shows the $Log_2$ fold change during necroptosis (with respect to control) for each identified transcript
Column K shows the $p$ value for each identified transcript during necroptosis (with respect to control)
Column L shows the $p_{adjusted}$ value for each identified transcript during necroptosis (with respect to control)

**The second tab ("all necrop padj<0.05") contains all genes detected in control and necroptotic cells that have $p_{adjusted}$ value < 0.05 during necroptosis:**
Column A shows the ensemble gene ID for detected transcripts.
Columns B-D show the values of read depth of transcripts in control cells (n=3)
Columns E-G show the values of read depth of transcripts in necroptotic cells (n =3)
Column H shows the HUGO gene nomenclature (hgnc_symbol) for identified transcript
Column I shows the description of hgnc_symbol
Column J shows the $Log_2$ fold change during necroptosis (with respect to control) for each identified transcript
Column K shows the $p$ value for each identified transcript during necroptosis (with respect to control)
Column L shows the $p_{adjusted}$ value for each identified transcript during necroptosis (with respect to control)

**Table S2 (Excel file). Results of transcriptomic data. Lipid related transcripts changing significantly in necroptotic cells compared to control are reported.**
**This table is provided as a separate spread sheet.**

Column A shows the ensemble gene ID for detected transcripts.
Columns B-D show the values of read depth of transcripts in control cells (n= 3)
Columns E-G show the values of read depth of transcripts in necroptotic cells (n= 3)
Column H shows the HUGO gene nomenclature (hgnc_symbol) for identified transcript
Column I shows the description of hgnc_symbol
Column J shows the $Log_2$ fold change during necroptosis (with respect to control) for each identified transcript
Column K shows the *p* value for each identified transcript during necroptosis (with respect to control)
Column L shows the $p_{adjusted}$ value for each identified transcript during necroptosis (with respect to control)

| Gene | Primer | Sequence | Identifier |
|---|---|---|---|
| *ELOVL7* | sense | 5’-GCA ATC CTC CAT GAA AAA GAA CT-3’ | Hs.PT.58.40 27298 |
| | antisense | 5’-CCA GCC TAC CAG AAG TAT TTG TG-3’ | |
| *FASN* | sense | 5’-GCA GTT CAC GGA CAT GGA-3’ | Hs.PT.58.39 640679 |
| | antisense | 5’-CTG GTG GCT CTT GAT GAT CAG-3’ | |
| *MLYCD* | sense | 5'-GAT GGA ATA AAA GAT CGC AGC A-3' | Hs.PT.58.39 208007 |
| | antisense | 5'-TTT GCA CGT GGC ACT GA-3' | |
| *HMGCR* | sense | 5'-CTG ACA TGC AGC CAA AGC-3' | Hs.PT.58.41 105492 |
| | antisense | 5'-GTT TAC CCT CGA TGC TCT TGT-3' | |
| *HMGCS1* | sense | 5'-GCC TTC TCC ACA TCT CTA TCA AAG-3' | Hs.PT.58.27 368554 |
| | antisense | 5'-CTC GGA TGT TGC TGA ATG ACT-3' | |
| *MCEE* | sense | 5'-ATG TGC TCC TAT TTT GAC CTC T-3' | Hs.PT.58.29 57789 |
| | antisense | 5'-GCA GAA AAA CAA GGC TGG AG-3' | |
| *PPARD* | sense | 5'-GCC ACT GTG TGA GTA TCA CG-3' | Hs.PT.58.38 472006 |
| | antisense | 5'-GGG AAA AGT TTT GGC AGG AG-3' | |
| *LIAS* | sense | 5'-TTG TCC TAA AGT CAA GCA GTC T-3' | Hs.PT.58.19 178010 |
| | antisense | 5'-CGT GTA CTG AAA CAT GCC AAG-3' | |
| *OXSM* | sense | 5'-GCA GCA GCC ATA CAC CTT A-3' | Hs.PT.58.38 451856 |
| | antisense | 5'-AGT ATC CAC AGC CTG TAC CA-3' | |
| *INSIG1* | sense | 5'-AAC GAT CAA ATG TCC ACC AAA G-3' | Hs.PT.58.25 883577 |
| | antisense | 5'-CAT TAA CCA CGC CAG TGC T-3' | |
| *ACACA* | *sense*<br>*antisense* | 5’-GTA CAT CGC TGA CAC TAG CTA C-3’<br>5’-CTG CCC ACA TCT CAT CCA AA-3’ | Hs.PT.56a.5 13712.g |
| *SCAP* | *sense*<br>*antisense* | 5’-CAC GAC AGA AAG AGA CAG AA-3’<br>5’-GAG AGC TGG TCC ATC ATG AAG-3’ | Hs.PT.58.45 442299 |
| *HPRT1* | sense | 5’-TTG TTG TAG GAT ATG CCC TTG A-3’ | Hs.PT.58v.4 5621572 |
| | antisense | 5’-GCG ATG TCA ATA GGA CTC CAG-3’ | |
| *GAPDH* | *sense*<br>*antisense* | 5’-ACA TCG CTC AGA CAC CAT G-3’<br>5’-TGT AGT TGA GGT CAA TGA AGG G-3’ | Hs.PT.39a.2 2214836 |

**Table S3.** Sequences of primers used for droplet digital PCR.

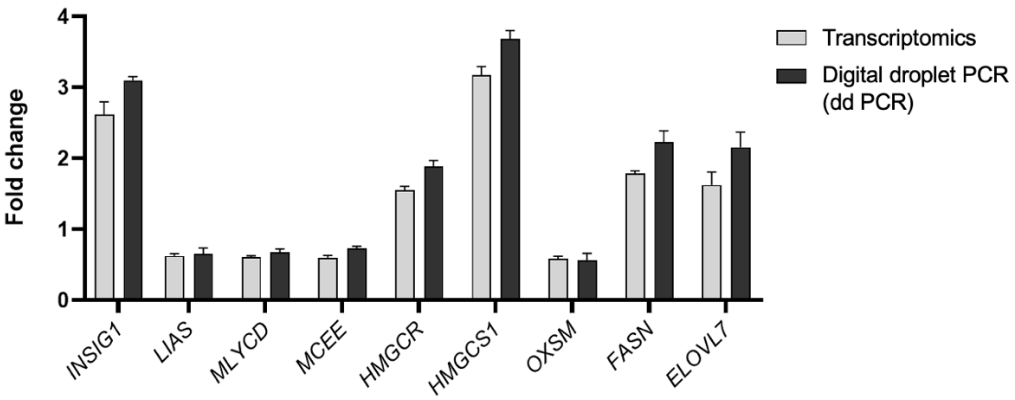


**Figure S1.** Validation of transcriptomics results via comparison of fold changes obtained from transcriptomics dataset and from independently measured samples by digital PCR experiments. Data represents mean ± 1 SD; n= 3. Gene abbreviations: insulin induced gene 1, *INSIG1*; Lipoic Acid Synthetase, *LIAS*; Malonyl-CoA Decarboxylase, *MLYCD*; Methylmalonyl-CoA Epimerase, *MCEE*; 3-hydroxy-3-methylglutaryl-CoA reductase, *HMGCR*; 3-hydroxy-3-methylglutaryl-CoA synthase 1, *HMGCS1*; squalene epoxidase, *SQLE*; 3-Oxoacyl-ACP Synthase, Mitochondrial, *OXSM*; fatty acid synthase, *FASN*; *ELOVL7*, Elongation of Very Long Chain Fatty Acids Protein 7.

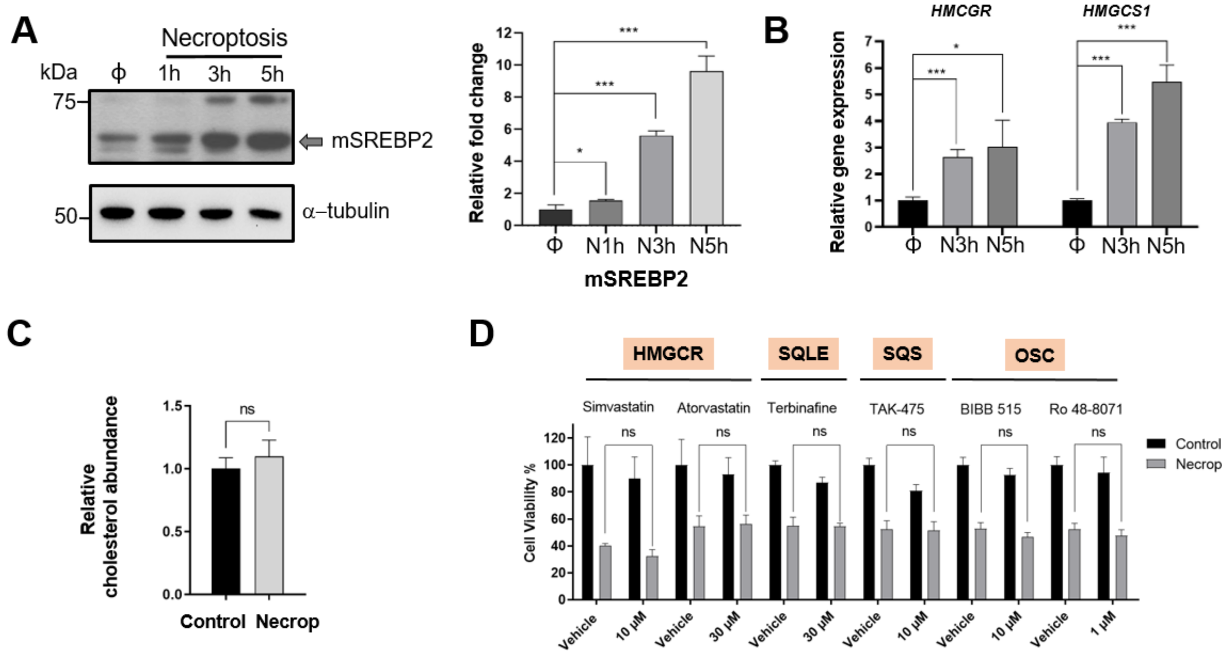


**Figure S2.** Potential SREBP2 involvement in necroptosis. (**A**) Western blot analysis for the time-dependent increase in SREBP2 activation during necroptosis. Cells were induced with necroptosis for 1 h, 3 h, and 5 h. The whole lysate samples were prepared and analyzed. Mature SREBP2 (mSREBP2) also increases time-dependently compared to control during necroptosis just like SREBP1. The right panel shows the quantification of mSREBP2 band intensities. There is a significant increase in activation during necroptosis compared to control cells. Data represent mean ± 1 SD; n= 3. * represents $p < 0.05$, *** represents $p < 0.001$. Φ represents DMSO control, N1h represents 1h necroptosis, N3h represents 3h necroptosis, N5h represents 5h necroptosis (**B**) SREBP2 target genes are upregulated during necroptosis. Fold changes in expression of *HMGCR* and *HMGCS1*, are calculated as the ratio of relative expression of each gene compared with *HPRT1* in necroptotic and control cells. Φ represents DMSO control, N3h represents 3h necroptosis, N5h represents 5h necroptosis. Data represent mean ± 1 SD; n= 3. * represents $p < 0.05$, *** represents $p < 0.001$. (**C**) LC-MS showed cholesterol level during necroptosis does not change. Data represent mean ± 1 SD; n= 3. ns represents $p > 0.05$. (**D**) Targeting the cholesterol biosynthesis pathway using small molecule inhibitors did not affect the necroptosis phenotype. HT-29 cells were plated and pretreated either with simvastatin, terbinafine, TAK-475, BIBB 515, or Ro 48-8071 for 24 h, atorvastatin for 48 h. After pretreatment cells were induced with necroptosis for 3h and subjected to an MTT cell viability assay. Data represent mean ± 1 SD; n ≤ 5, ns represents $p > 0.05$. Enzyme abbreviations: 3-hydroxy-3-methylglutaryl-CoA reductase, HMGCCR; squalene epoxidase, SQLE; 3-Oxoacyl-ACP synthase; squalene synthase, SQS; 2,3-oxidosqualene cyclase, OSC.

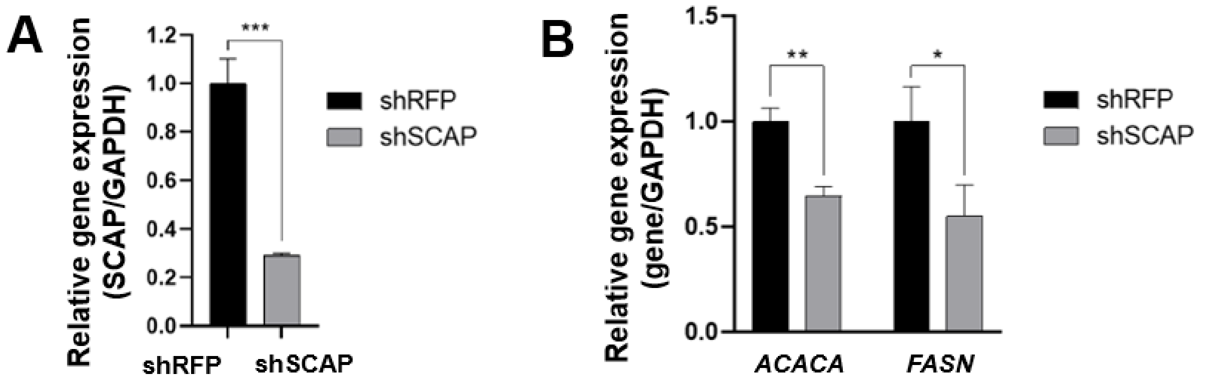


**Figure S3.** Knockdown models affecting SREBP1 activation. (**A**) Transfection efficiency of SCAP in HT-29 cells. Red fluorescent protein (RFP) was used as a knockdown control. Fold changes in expression of *SCAP* are calculated as the ratio of relative expression of SCAP compared with *GAPDH* in necroptotic and control cells. Data represent mean ± 1 SD; n = 3. *** represents $p < 0.001$. (**B**) Depletion of SREBP1 target genes in shSCAP cells. *ACACA* and *FASN* are downregulated in shSCAP knockdown cells. Fold changes in expression of *ACACA* and *FASN* are calculated as the ratio of relative expression of the target genes compared with *GAPDH* in necroptotic and control cells. Data represent mean ± 1 SD; n = 3. * represents $p < 0.05$, ** represents $p < 0.01$.

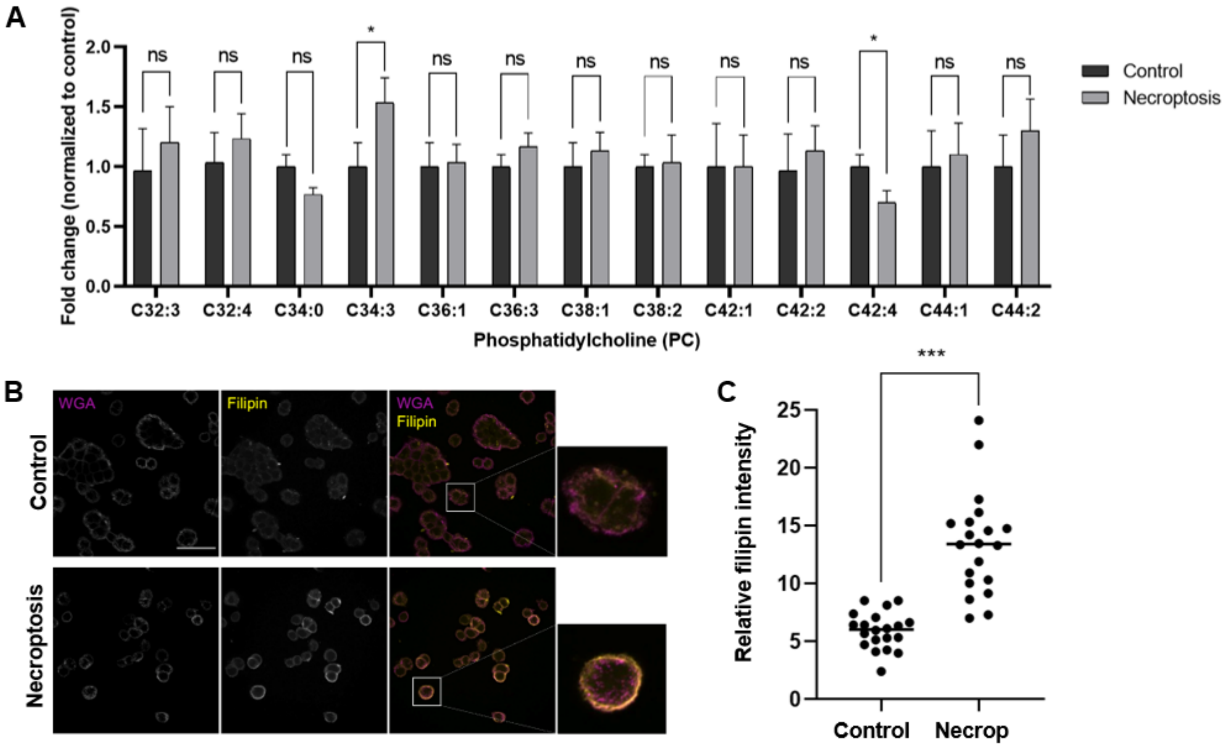


**Figure S4.** (**A**). Phosphatidylcholine (PC) levels during necroptosis. The relative abundance of PC species (normalized to control) showed no significant changes in necroptotic cells compared to control cells alone. Data represents mean ± 1 SD; n= 3, ns represents not significant, * represents $p < 0.05$. (**B**) Fluorescence microscopy images showed cholesterol localization at the plasma membrane during necroptosis. Cells were stained with filipin which binds to free cholesterol. Representative images from at least three experiments are shown. To illustrate changes in the plasma membrane more clearly, magnified images are shown next to the merged images. The white scale bar represents 50 µm. (**C**) Quantitative analysis of cholesterol localization during necroptosis. One representative image from the control or necroptotic condition was selected for quantification (n= 20, *** represents $p < 0.001$, see supporting information method details for the quantification of filipin).
